# Widespread distribution and evolution of poxviral entry-fusion complex proteins in giant viruses

**DOI:** 10.1101/2023.01.19.524718

**Authors:** Sheng Kao, Chi-Fei Kao, Wen Chang, Chuan Ku

## Abstract

Poxviruses are known to encode a set of proteins that form an entry-fusion complex (EFC) to mediate virus entry. However, the diversity, evolution, and origin of these EFC proteins remain poorly understood. Here we identify the EFC protein homologs in poxviruses and other giant viruses of phylum *Nucleocytoviricota*. The 11 EFC genes are present in almost all poxviruses, with the two smallest, G3 and O3, absent in *Entomopoxvirinae* and basal lineages of *Chordopoxvirinae*. Five of the EFC genes are further grouped into two families, A16/G9/J5 and F9/L1, which are widely distributed across other major lineages of *Nucleocytoviricota*, including metagenome-assembled genomes, but are generally absent in viruses infecting algae or non-amoebozoan heterotrophic protists. The A16/G9/J5 and F9/L1 families co-occur, mostly as single copies, in 93% of the non-*Poxviridae* giant viruses that have at least one of them. Distribution and phylogenetic patterns suggest that both families originated in the ancestor of *Nucleocytoviricota*. In addition to the *Poxviridae* genes, homologs from each of the other *Nucleocytoviricota* families are largely clustered together, suggesting their ancient presence and vertical inheritance. Despite deep sequence divergences, we observed noticeable conservation of cysteine residues and predicted structures between EFC proteins of *Poxviridae* and other families. Overall, our study reveals widespread distribution of these EFC protein homologs beyond poxviruses, implies the existence of a conserved membrane fusion mechanism, and sheds light on host range and ancient evolution of *Nucleocytoviricota*.

**Importance:** Fusion between virus and host membranes is critical for viruses to release genetic materials and to initiate infection. Whereas most viruses use a single protein for membrane fusion, poxviruses employ a multi-protein entry-fusion complex (EFC). We report that two major families of the EFC proteins are widely distributed within the virus phylum *Nucleocytoviricota*, which include poxviruses and other dsDNA “giant viruses” that infect animals, amoebozoans, algae, and various microbial eukaryotes. Each of these two protein families is structurally conserved, traces its origin to the root of *Nucleocytoviricota*, was passed down to the major subclades of *Nucleocytoviricota*, and is retained in most giant viruses known to infect animals and amoebozoans. The EFC proteins therefore represent a potential mechanism for virus entry in diverse giant viruses. We hypothesize that they may have facilitated the infection of an animal/amoebozoan-like host by the last *Nucleocytoviricota* common ancestor.

## Introduction

Viral infection begins with entry into the host cell. As a prototypic member of *Poxviridae*, vaccinia mature virus binds to cell surface glycosaminoglycans and laminin (1–3) and induces actin-dependent endocytosis of virus particles (4–7). Then the endosomal low pH triggers membrane fusion between viral membrane and endosomal membrane, releasing vial core into cytoplasm (8). For enveloped viruses such as poxviruses, the timing of fusion activation is controlled by viral fusion proteins that are activated by acidic environment. To date most viruses use a single viral protein to trigger fusion with host membrane (9). However, vaccinia virus (VacV) contains an entry-fusion protein complex (EFC) of 11 components, A16, A21, A28, F9, G3, G9, H2, J5, L1, L5, and O3, to execute membrane fusion during virus entry ((10, 11) and references therein). Deletion of individual components of the EFC generated VacV particles with low infectivity, but how EFC mediates viral membrane fusion remains unknown (11).

*Poxviridae* belongs to the recently established phylum *Nucleocytoviricota* (12). They are nucleocytoplasmic large dsDNA viruses that infect all major lineages of eukaryotes, from animals and algae to amoebae and other microbes (13), and are commonly referred to as giant viruses (GVs) (14–17) for having genomes and virions comparable in size to small bacteria. GVs encode diverse proteins rarely or never found in other viruses, with very few genes widely shared across major GV lineages and most genes acquired in a lineage-specific manner at the family or lower taxonomic level (18). Virus-cell interactions, including the processes and mechanisms of cell entry, are poorly understood for most GVs. Similar to poxviruses, other GVs mostly have at least one outer envelope membrane, internal membrane, or both. In addition to animal viruses, membrane fusion has been observed at the initial stage of infection by GVs of algae (e.g., chloroviruses (19)) and other microbial eukaryotes. In amoeba-infecting GVs, infection typically begins with phagocytosis followed by capsid opening and fusion of the internal membrane with the phagosome (17, 20). The low endosomal pH has also been suggested to induce membrane fusion for different GVs (21), including those infecting vertebrates (22, 23) and amoebozoans (24). Recent studies on African swine fever virus (ASFV) found that its pE248R (25) and pE199L (26) proteins are required for viral membrane fusion and are distant homologs of VacV EFC proteins A16/G9/J5 and F9/L1, respectively. Homologs of these proteins have also been detected in some other GVs through BLAST searches (26) and gene clustering (27), but their distribution across all the GVs, including numerous recently reported metagenome-assembled genomes (MAGs) (17, 28, 29), remains to be determined. It is also unclear whether the poxviral EFC homologs may function as an evolutionarily conserved fusion machinery in other GVs.

To better understand GV gene repertoires and virus entry, this study aims to unravel the distribution patterns, evolutionary history, and conservation of EFC proteins in poxviruses and the other GVs. Through phylogenetic analyses and sequence comparisons, our results provide insights into the origin, diversification, and duplications of EFC proteins in GVs. Furthermore, predicted models shed light on the protein structural similarity across divergent GV lineages. These findings suggest an important functional role of poxviral EFC proteins in other GVs.

## Materials and Methods

### Identification of EFC homologs

To identify genes across giant viruses that encode homologs of EFC proteins, we searched for orthogroups (gene clusters) containing individual VacV EFC genes in the all-against-all gene clustering dataset of a recent study on gene contents of 207 GV genomes (18), including 51 from *Poxviridae*. In addition to these genomes, we added 4 genomes covering underrepresented lineages of Poxviridae, including carp edema virus strain FTI2020 (30), saltwater crocodilepox virus subtype 1 (31), teiidaepox virus 1 (32), and cheloniid poxvirus 1 (33). The presence-absence patterns of the EFC genes in all poxviruses were verified through literature search and using TBLASTN (BLAST v2.6.0 (34)) with VacV EFC proteins against the other poxvirus genome nucleotide sequences. For non-poxvirus genomes, EFC gene homologs in *Asfarviridae, Iridoviridae, Mimiviridae*, and *Pithoviridae* reported previously (26, 35) were included and used to find other EFC homologs that were clustered together in the same orthogroups. Furthermore, we used these EFC homologs encoded by the predominantly cultured viruses as queries and performed a BLASTP (34) search against predicted proteins in two datasets of MAGs of environmental giant viruses sampled from diverse ecosystems (28, 29) (query coverage ≥ 30% and e-value ≤ 1 × 10^−5^). A similar BLASTP search (query coverage ≥ 30%, alignment length ≥100, and e-value ≤ 1 × 10^−5^) against the NCBI non-redundant protein database (36) was used to detect any additional EFC homologs in genome sequences of other giant viruses, other viruses and cellular organisms (excl. hits to highly repetitive sequences or parts absent in poxvirus homologs).

### Phylogenetic analyses

The identified sequences in each of the two protein families, A16/G9/J5 and F9/L1 (Table S1), were aligned using MAFFT v7.310 (37) with the default parameters. Maximum likelihood trees were constructed from the alignments using IQ-TREE v2.2.0 (38) with the best-fit model selected using ModelFinder (39). The trees were visualized using Figtree v1.4.4 and the package ggtree v3.4.4 (40) in RStudio (2022.02.3 build 492).

### Structural modeling

The VacV F9 (PDB: 6CJ6) (41) and L1 (PDB: 1YPY) (42) protein structures were previously determined by X-ray crystallography. We predicted the structure of F9/L1 proteins of selected GVs using AlphaFold2 (AF2) web server v1.4 (https://colab.research.google.com/github/sokrypton/ColabFold/blob/main/AlphaFold2.ipynb; accessed in Oct. 2022) with the default settings without AMBER relaxation or PDB templates (43, 44). All sequence alignments and templates were obtained by MMseqs2 (45). In order to evaluate the model quality, the predicted Local Distance Difference Test (pLDDT) score of each amino acid (aa) was calculated. Only models having both the mean pLDDT score and half of the residue pLDDT scores above 70 were retained for further analyses. The highest-scored model of each F9/L1 protein was superimposed with the VacV L1 protein structure using PyMOL Molecular Graphics System v1.8 (46). Protein structure predictions of A16/G9/J5 homologs were also performed using AF2 with the same parameters and criteria. Because the high sequence divergence and length variation, our structural comparisons focused on the conserved N-terminal ectodomains of F9/L1 (VacV L1 aa 1–175) and A16/G9/J5 (VacV J5 aa 1–89), which contain characteristic disulfide bonds suitable for protein superposition comparison. The root-mean-square deviation (RMSD) value was calculated in PDBeFold using the secondary-structure matching (SSM) tool (47). All the protein structural and modeling drawings were prepared using PyMOL as described in the user manual.

## Results

### Presence-absence of EFC genes

Based on a recent gene clustering dataset of 207 GVs (18), mostly isolated and cultured viruses with complete or nearly complete genome sequences, we detected homologs of EFC genes in 124 GVs (Figure 1). The 11 EFC genes are divided into 8 gene families. One gene family contains VacV A16/G9/J5 and another F9/L1, which correspond to NCVOG1122 and NCVOG0211 (27), respectively. Most EFC genes, including A16, G9, J5, F9, L1, H2, A21, and A28, are present as a single copy in all poxviruses (except for two duplicated F9 copies in the pseudocowpox virus; Figure 1 and Table S1). TBLASTN searches suggest that the two smallest EFC proteins, G3 and O3, are present in core *Chordopoxvirinae* but absent in salmon gill poxvirus, and Nile crocodilepox virus, and *Entomopoxvirinae*. L5 is present in all poxviruses except salmon gill poxvirus. For the four additional poxviruses included in this study, we found that carp edema virus and saltwater crocodilepox virus have the same presence-absence patterns as salmon gill poxvirus and Nile crocodilepox virus, respectively. Similar to the avipoxviruses they are the most closely related to, teiidaepox virus and cheloniid poxvirus have all the EFC genes (Table S2). Based on the core gene tree as the backbone phylogeny of GVs (Figure 1), G3 and O3 likely originated in the common ancestor of core *Chordopoxvirinae* after its divergence from crocodilepox virus. H2, A21, A28, and L5 all trace their origins to the root of *Poxviridae*, with L5 lost during the evolution of the salmon gill poxvirus lineage.

**Figure 1.**
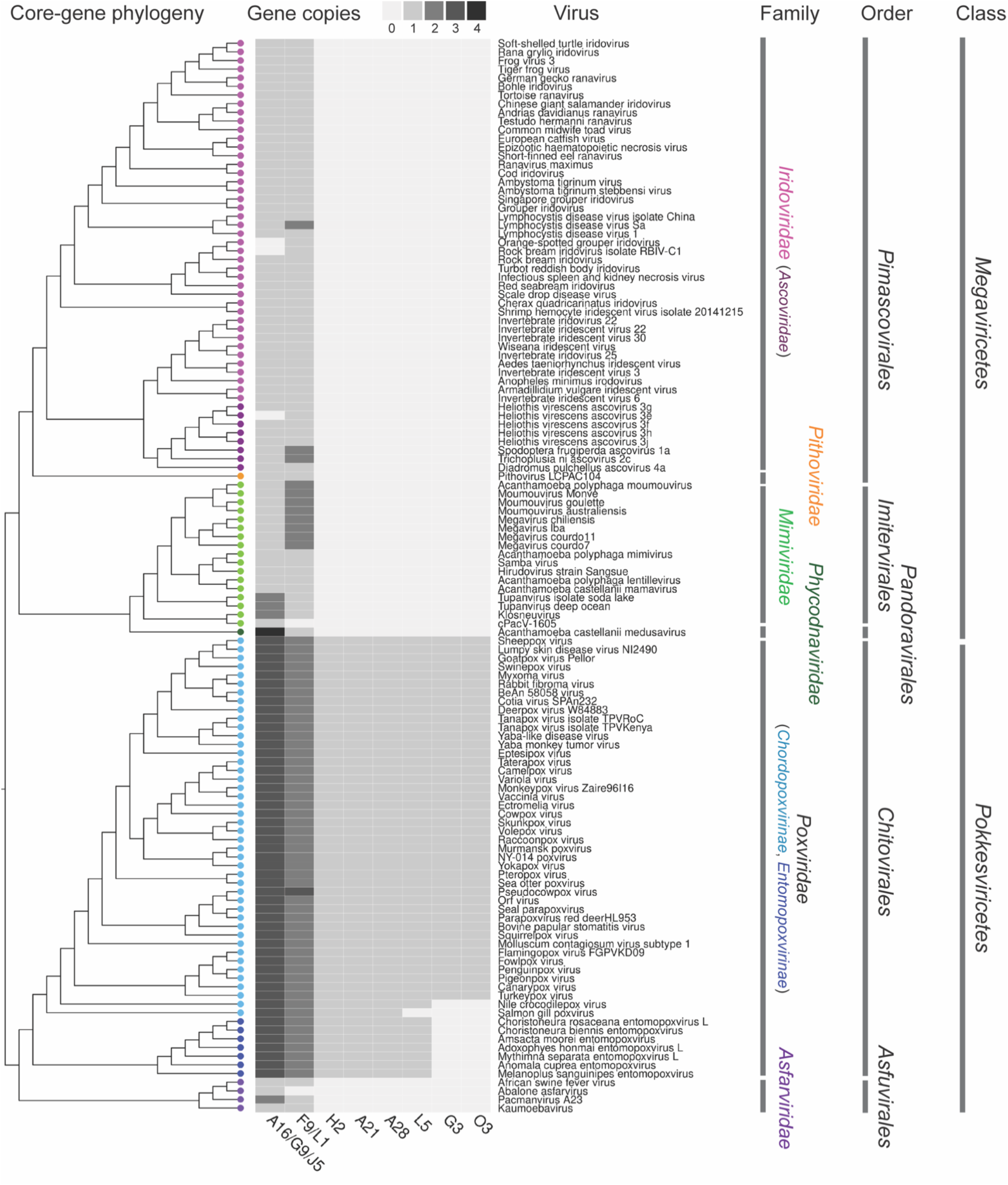
Distribution and copy numbers of entry-fusion complex (EFC) gene homologs in *Poxviridae* and other *Nucleocytoviricota* families. The 11 EFC proteins in vaccinia virus are grouped into 8 families, including one that includes A16, G9, and J5 and another that combines F9 and L1. Out of a total of 207 viruses included in a recent study (18), 124 have at least one EFC homolog and their phylogenetic relationships based on five core proteins (18) are shown on the left. The family-level (18), order-level (48), and class-level (12) taxa are indicated on the right.

Homologs of A16/G9/J5 and F9/L1 are present in 124 out of the 207 GV genomes (Figure 1). The viruses belong to both classes of *Nucleocytoviricota*, covering 6 of the 7 major family-level clades as defined previously (18) and 5 of the 6 order-level clades delineated in another study (48). Given the broad distribution across major GV lineages, it is notable that none of the isolated GVs known to infect algae have any of the EFC genes. Where A16/G9/J5 and F9/L1 genes are present, they generally co-occur, as in 93.2% (68/73) of the non-*Poxviridae* GV genomes that have at least one of them (Figure 1). Unlike in *Poxviridae*, both gene families mostly have only one single copy in the non-*Poxviridae* genomes, with some exceptions being medusavirus (4 genes of A16/G9/J5) and moumouviruses/megaviruses (2 genes of F9/L1).

In addition to the 207 genomes of mostly isolated GVs, our search for EFC homologs extended to other GV genomes (Table S1). These include viruses in the newly proposed *Mininucleoviridae*, which have the smallest genomes (70-74 kb) in *Nucleocytoviricota* (49), *Marseilleviridae* genomes not included in the 207-genome dataset, and various MAGs (28, 29) that have been tentatively assigned to previously proposed *Nucleocytoviricota* families or newly delineated higher-level groups called “super-clades” (29). Although the MAGs tend to be more fragmented and partial genome assemblies, many of them have both A16/G9/J5 and F9/L1 genes detected in their genomes (Table S1).

### Sequence conservation and phylogenies of EFC genes

As in poxviruses, A16/G9/J5 and F9/L1 proteins in other GVs are cysteine-rich and have predicted transmembrane domains at or near the C-terminal end (Figures 2A and 3A). Despite sequence length variation, a conserved region is found in all A16/G9/J5 proteins, including the shortest protein J5 in poxviruses. Similarly, a conserved region is also found in all F9/L1 proteins.

**Figure 2.**
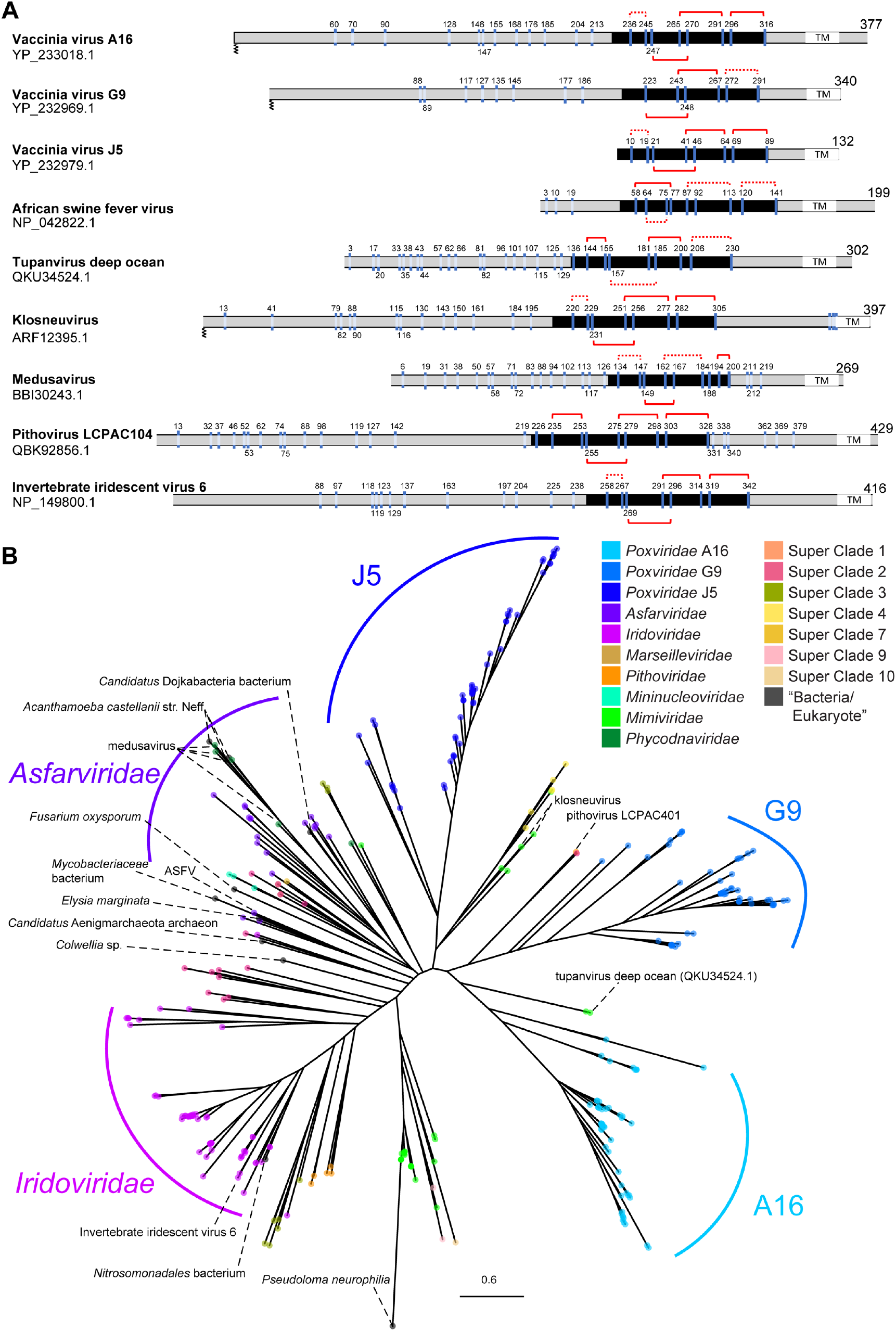
Positions of cysteine residues in representative homologs and phylogeny of the A16/G9/J5 gene family. (**A**) Cysteine residues are marked and numbered for representative sequences from different families. Disulfide bonds in the conserved region (black; J5-like domain) are denoted by red solid (found in structures predicted by AlphaFold2) or dotted (possible but not predicted) connections. Transmembrane domains (TM) and known or predicted N-terminal myristoylation are indicated. (**B**) Maximum-likelihood phylogenetic tree of A16/G9/J5 protein sequences from cultured isolates and metagenome-assembled genomes of giant viruses and the nr database of NCBI. ASFV: African swine fever virus.

**Figure 3.**
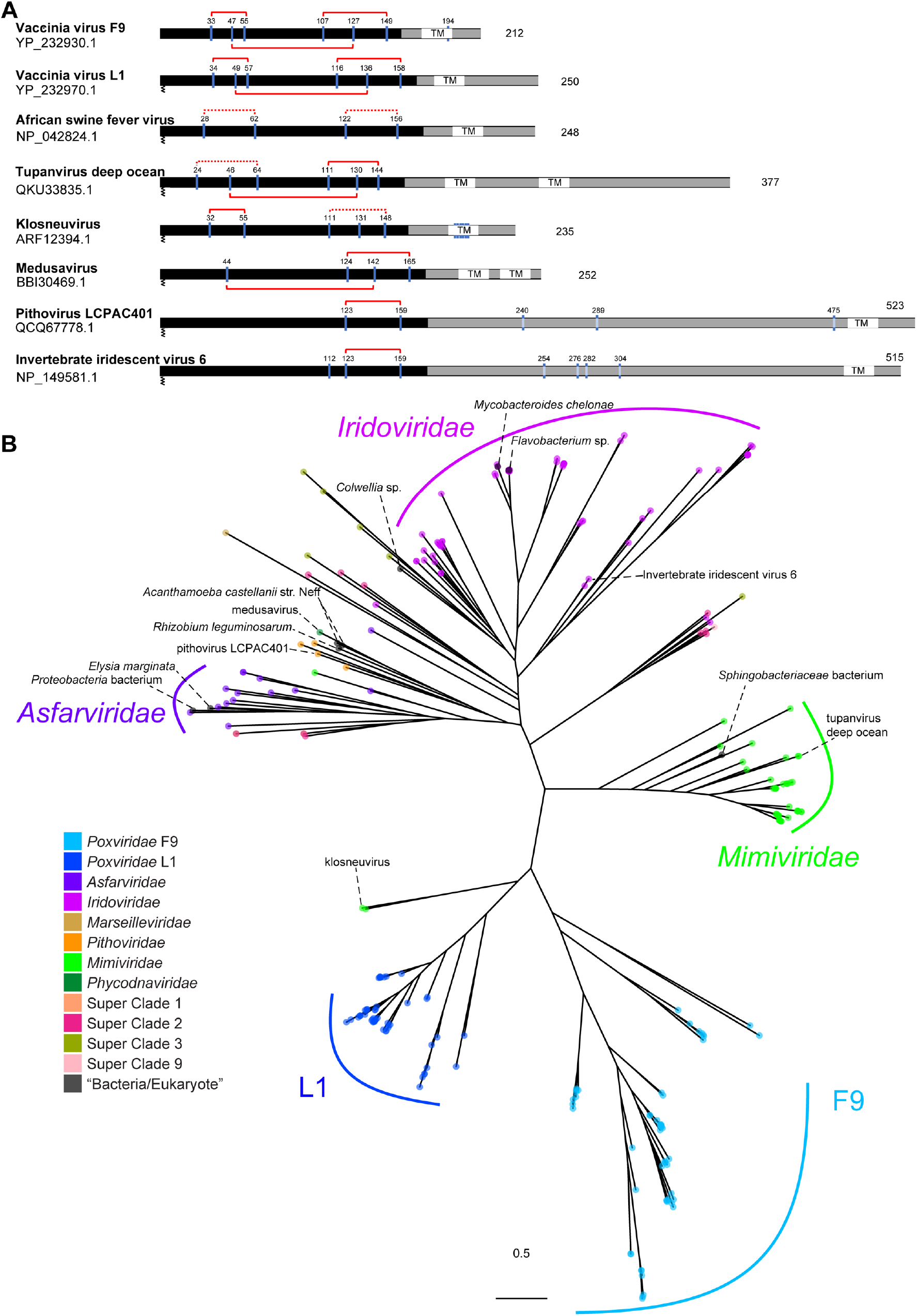
Positions of cysteine residues in representative homologs and phylogeny of the F9/L1 gene family. (**A**) Cysteine residues are marked and numbered for representative sequences from different families. Disulfide bonds in the conserved region (black) are denoted by red solid (found in structures predicted by AlphaFold2 or based on X-ray crystallography) or dotted (possible but not predicted). Transmembrane domains (TM) and known or predicted N-terminal myristoylation are indicated. (**B**) Maximum-likelihood phylogenetic tree of A16/G9/J5 protein sequences from cultured isolates and metagenome-assembled genomes of giant viruses and the nr database of NCBI.

Maximum likelihood phylogenies (Figures 2B and 3B; Data S1 and S2) were infered for both gene families from all the sequences listed in this study (Table S1). All five genes in *Poxviridae* (i.e., A16, G9, J5, F9, and L1) each form a monophyletic group, suggesting each of these genes has a single origin, was present in the last common ancestor of *Poxviridae*, and since then has not been transferred between poxviruses and any other genomes based on the current datasets. Within both A16/G9/J5 (Figure 2B) and F9/L1 (Figure 3B) trees, the closer relationships between different *Poxviridae* genes also implies that they likely share a common origin to the exclusion of most non-*Poxviridae* homologs. Homologs of other GV families, including *Iridoviridae* and *Asfarviridae*, form largely monophyletic groups in both trees. *Mimiviridae* forms one major clade in both trees, except for some homologs found in klosneuviruses and tupanviruses that are more closely related to poxvirus genes. These phylogenetic patterns indicate that after the divergence of A16, G9, and J5 (Figure 2B) or F9 and L1 (Figure 3B) and before the last *Poxviridae* common ancestor, some transfer events took place between early poxvirus ancestors and other genomes that eventually led to the presence of these poxvirus-related homologs in some mimiviruses, superclade 4 MAGs, and pithovirus LCPAC401 (Figures 2B and 3B).

A16/G9/J5 and F9/L1 homologs were also detected in non-GV genomes based on the BLASTP search against nr (Table S1). These include *Acanthamoeba castellanii* str. Neff, which has been widely used as a lab host for GV isolation and cultivation. The close relatioships between medusavirus and *A. castellanii* str. Neff in both gene family trees are consistent with the report of high gene sharing between these two genome sequences (50). The other genome sequences come from eukaryotes (*Elysia marginata, Fusarium oxysporum, Pseudoloma neurophilia*), one archeon (*Candidatus* Aenigmarchaeota archaeon), and a few bacteria. In total, these 11 A16/G9/J5 and 9 F9/L1 sequences from non-GV genomes are relatively few compared with the ones detected in GVs, and they tend to be found in nested positions within clades formed by GV sequences. Some of the non-GV genomes are incomplete assemblies, with the EFC genes found on small contigs. For example, the *Colwellia* sp. is a MAG from oxic subseafloor aquifer (51) with 422 contigs (JAESPX010000000), where A16/G9/J5 and F9/L1 genes are located on a 9732-bp contig (Table S1). Therefore, we cannot rule out that some of the EFC genes might actually be contaminating sequences from GVs, which are widely distributed in various ecosystems. Given the overall patterns of distribution and phylogenetic relationships, the A16/G9/J5 and F9/L1 protein families likely originated in *Nucleocytoviricota* or was transferred to a *Nucleocytoviricota* ancestor from some other lineage that has not been sampled or has gone extinct. Since these proteins are present in major lineages of both primary branches (classes *Pokkesviricetes* and *Megaviricetes*; Figure 1) of *Nucleocytoviricota*, their presence can be traced back to the last *Nucleocytoviricota* common ancestor (LNCA). After LNCA diversified into descendant lineages, A16/G9/J5 and F9/L1 genes duplicated in the ancestor of *Poxviridae* and have been maintained as three and two copies, respectively, throughout *Poxviridae* evolution.

### Genomic locations

In addition to co-occurrence of A16/G9/J5 and F9/L1 in most genomes, we observe some interesting patterns of their genomic locations (Figure 4). A16/G9/J5 and F9/L1 genes are often in close proximity (next to each other or within a few genes), such as those in ASFV, kaumoebavirus, and klosneuvirus (Figure 4A). The *Asfarviridae* GVs, ASFV and kaumoebavirus, both have a small ORF between their A16/G9/J5 and F9/L1 genes that run in opposite directions. Similarly in the MAG *Colwellia* sp., the A16/G9/J5 and F9/L1 copies are neighboring genes in opposite directions. Four copies of A16/G9/J5 occur within a region of 20 genes in the medusavirus genome. In a major subgroup of *Mimiviridae*, moumouviruses (lineage B), megaviruses (lineage C), and cotonvirus all have two adjacent copies of F9/L1, but there is only one copy in mimiviruses/mamaviruses (lineage A) (Figure 4B). Based on the F9/L1 phylogeny (Data S2) and the phylogeny of these viruses (52), a tandem duplication of the ancestral F9/L1 gene occurred in the common ancestor of these viruses, with both copies preserved in all lineages except mimiviruses/mamaviruses, which retained only the upstream copy of the two.

**Figure 4.**
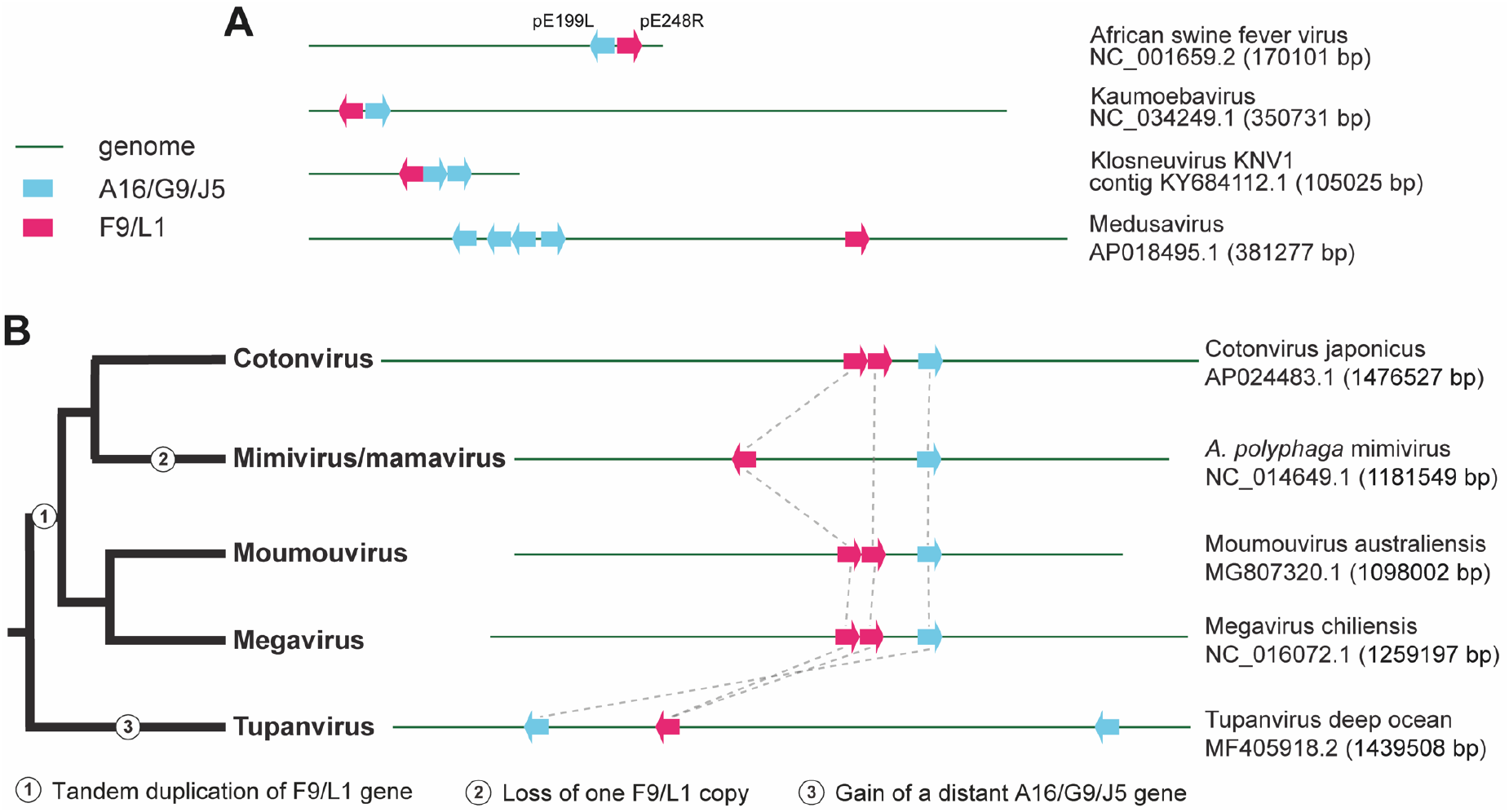
Genomic locations of A16/G9/J5 and F9/L1 genes in selected GV genomes. (**A**) Close proximity of both A16/G9/J5 and F9/L1 genes or multiple A16/G9/J5 genes. (**B**) Evolutionary dynamics of F9/L1 and A16/G9/J5 genes in a major clade of *Mimiviridae*. Dashed lines indicate orthology. Gene size and genome size are not drawn to scale.

### Structural conservation

To investigate whether the identified GV EFC homologs exhibit any conserved protein structure, we took advantage of the recently reported AI system AlphaFold2 for predicting protein structure with high accuracy (43, 44). We compared the disulfide bonding patterns of non-*Poxviridae* homologs with those of VacV proteins (Figure 2A and 3A) based on closely spaced cysteine-pairs that are either known to or can potentially form disulfide bonds in the structural models. Interestingly, we found that the disulfide bonding patterns represent an evolutionary conserved structural feature of EFC proteins.

The VacV L1 was the first EFC protein with a resolved atomic structure (42) (Figure 5). It comprises five α-helices and four β-strands (order: α1-β1-β2-α2-α3-α4-β3-β4-α5) forming a packed helical bundle juxtaposed to a pair of β-sheets (Figure 5A). Multiple sequence alignment revealed the presence of six conserved cysteines throughout the poxviral L1 orthologs. VacV L1 has a disulfide topology of 1-3_2-5_4-6 with disulfide bonding between Cys34 and Cys57, Cys49 and Cys136, and Cys116 and Cys158 (Figure 3A). More recently, the structure of the ectodomain of VacV F9 was also determined (41). Despite the low sequence identity (27%), superposition of L1 and F9 ectodomain revealed a well-aligned 3-dimensional conformation with an RMSD of 2.06 Å (Data S3). To validate the accuracy of AF2 prediction in EFC proteins, we also predicted the VacV L1 and F9 structures using AF2 and calculated the RMSD between the crystalized and the modeled structures upon superposition. The results showed that AF2 can accurately predict L1 and F9 structures with an RMSD of 1.75 Å across 138 Cα atoms and 2.30 Å across 143 Cα atoms, respectively.

**Figure 5.**
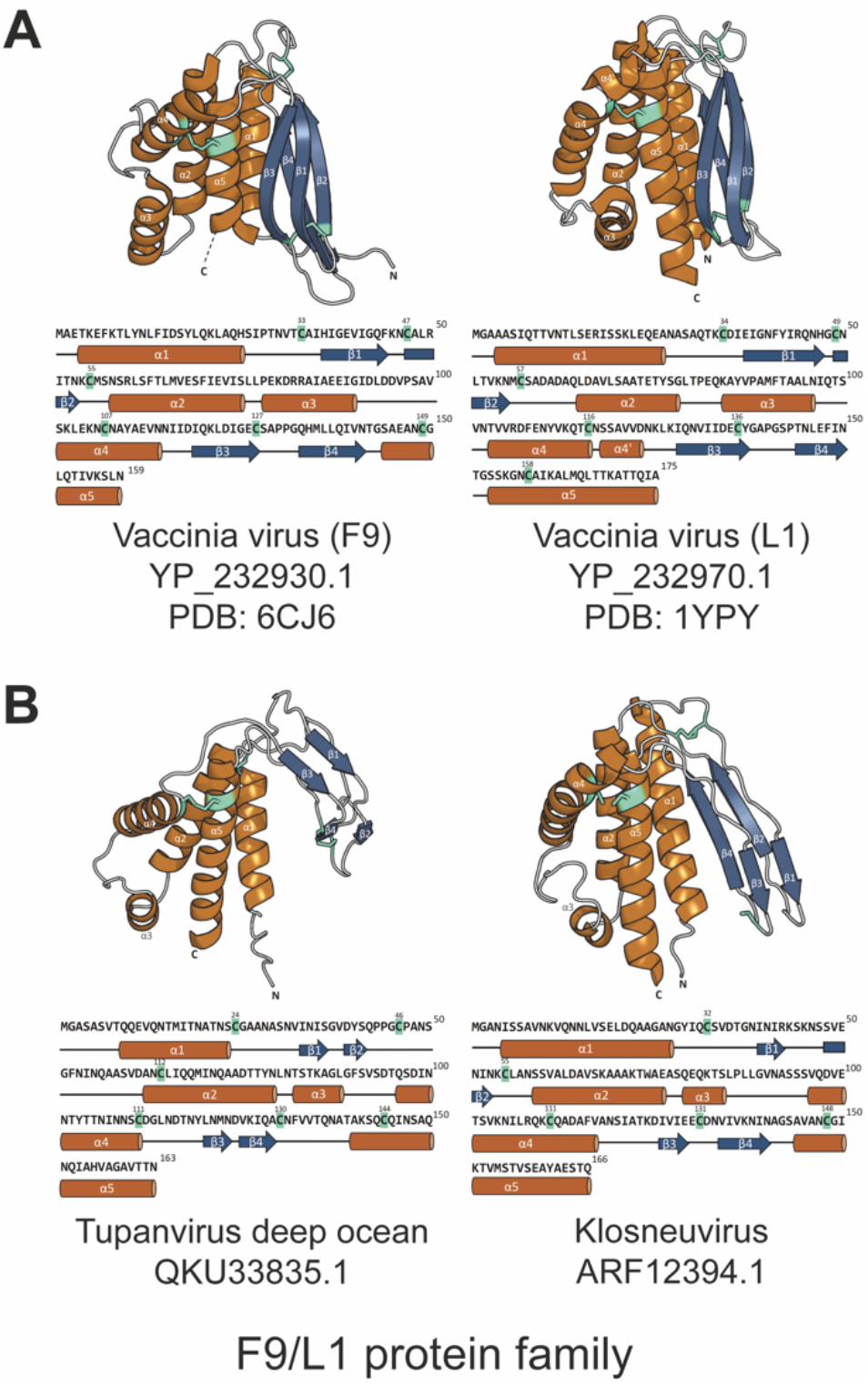
Overall structures and secondary structural elements of N-terminal ectodomain in F9/L1 family. (**A**) Cartoon representation of VacV F9 and L1 protein structures, which were determined by x-ray crystallography (41, 42). Both molecules consist of a five-helix bundle juxtaposing to two β-sheets. Three pairs of conserved disulfide bonds are found in F9 and L1. (**B**) Cartoon representation of AlphaFold2-predicted structures of N-terminal ectodomains of homologs in two other GVs. The modeled ectodomains, which comprise five α-helices packed against two β-sheets, exhibit structural similarity as those seen in F9 and L1. The relative positions of disulfide bonds (three pairs in QKU33835.1 and two pairs in ARF12394.1) are also similar, particularly the last pair bridging the helices α4 and α5. Cyangreen sticks indicate disulfide bonds. The α-helices and β-strands are labeled and colored orange and dark blue, respectively. The corresponding amino acid sequences annotated with secondary structural features are shown below the 3D models.

We then used AF2 to predict the protein structures of F9/L1 homologs in two non-*Poxviridae* GVs, including tupanvirus deep ocean (QKU33835.1) and klosneuvirus (ARF12394.1) (Figure 5B). High mean pLDDT scores (70.97 and 70.43, respectively) were obtained for their N-terminal ectodomains when compared with the VacV L1 and F9 ectodomains. The predicted protein structure of QKU33835.1 and ARF12394.1 both contain five helices and four β-strands in an order of α1-β1-β2-α2-α3-α4-β3-β4-α5, which is identical to that of the VacV L1 and F9. Despite some variation in local structures, the relative positions between the helical bundle and β-sheets/loops are the same. Furthermore, although the numbers of cysteine and the corresponding disulfide bonds vary, the structural modeling suggested that the last pair of cysteines connecting the helices α4 and α5 appears to be evolutionarily conserved in the F9/L1 family.

For the A16/G9/J5 family, we focused on the widely conserved structural domain (called J5-like domain hereafter) corresponding to the VacV J5 N-terminal ectodomain from aa 1 to 89 (Figure 2A). Although none of VacV A16, G9, and J5 protein structure has been resolved, the mean pLDDT score of the J5-like domain is all above 70 (VacV A16: 75.05; VacV G9: 79.89; VacV J5: 87.01). The AF2-predicted model of the J5 ectodomain contains six short α-helices (α1-6) connected by loops and a long C-terminal extended loop (aa 64 to 89) (Figure 6A). A total of eight cysteines likely form four disulfide bonds that sequentially connect the helices α1 and α2 (cys10 and cys19), helices α2 and α5 (cys21 and cys46), helices α4 and α6 (cys41 and cys64), and the C-terminal loop (cys69 and cys89) (Figure 6A). Both the structural features and the disulfide topology of 1-2_3-5_4-6_7-8 are well conserved in the corresponding domains in vaccinia A16 and G9 protein (Figure 6a), except that VacV G9 lacks the two cysteines forming the first pair of disulfide bond.

**Figure 6.**
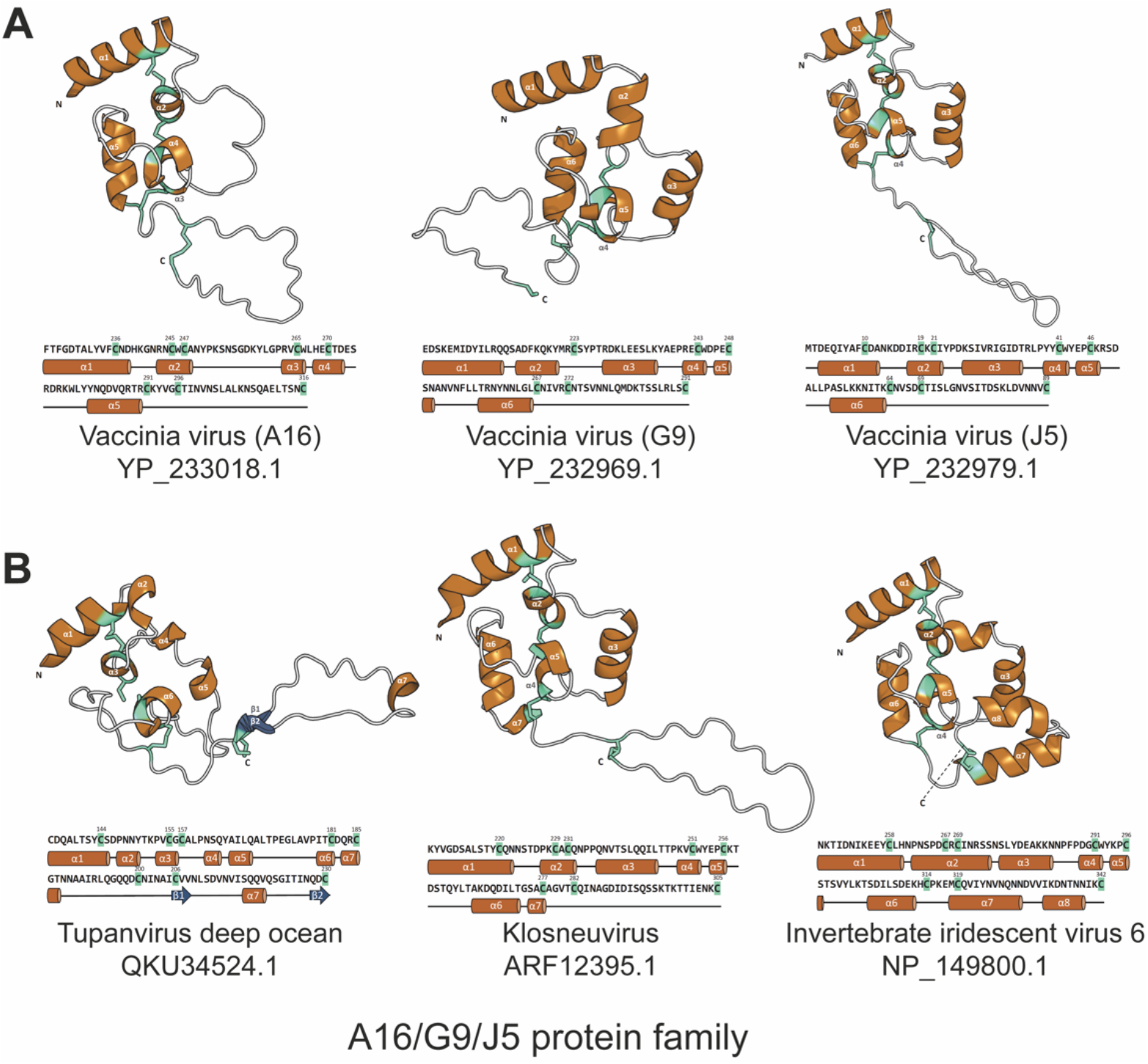
Overall structures and secondary structural elements of AlphaFold2-predicted J5-like domains in A16/G9/J5 family. (**A**) VacV A16, G9, and J5. (**B**) Homologs from other GVs. A common fold in the J5-like domain is comprised mainly by α-helices and a C-terminal extended loop. There are three to four pairs of conserved disulfide bonds (cyangreen sticks) which constrain the core architecture; however, the first pair that connects helices α1 and α2 is missing in G9. The α-helices and β-strands are labeled and colored orange and dark blue, respectively. The corresponding amino acid sequences are shown below each 3D model and annotated with the secondary structural features.

We then extended the structural prediction to three other GV homologs of A16/G9/J5 (Figure 6B). The mean pLDDT scores of the J5-like domain (Figure 2A) in these viral homologues are all above 70 (tupanvirus deep ocean QKU34524.1: 73.13, klosneuvirus ARF12395.1: 80.67; invertebrate iridescent virus 6 NP_149800.1: 86.43), supporting that these predictions are of high confidence and accuracy. These J5-like domains in non-*Poxviridae* GVs are similarly built by short 6-8 α-helices that are linked by loops (Figure 6B). Both the klosneuvirus and tupanvirus deep ocean homologs have an extended C-terminal loop. Besides, the cysteine spacing pattern also shows conserved distances between the second and third cysteines (2 residues) as well as the fourth and fifth cysteines (4 to 5 residues) that anchor the central helices and thus constrain the core region.

Besides the protein structural conservation described above, we also notice that VacV A16, G9 and L1 all have a conserved N-terminal myristoylation motif “MG” which anchors the protein in membrane and are of functional importance (53–55). The presence of the MG motif in most F9/L1 (Figure 3) proteins implies that post-translational lipid modification for N-terminus anchoring on membrane is well conserved in the F9/L1 family.

### Expression patterns and protein localization

The abundant protein-coding genes in GV genomes are typically expressed in transcriptional kinetic classes that are turned on and off sequentially (56–60). Through literature research, we found that EFC genes are mostly expressed late in the transcription programs of both *Poxviridae* and non-*Poxviridae* GVs. In a transcriptomic analysis of VacV, 7 EFC genes were expressed in the late kinetic class (A16, G9, L1, A21, A28 G3), and 1 in the early/late class (F9), whereas the expression levels of 4 genes were not well determined (J5, H2, L5, O3). In terms of localization, these 11 EFC proteins are all known to be embedded in the mature virion membrane of VacV (11). The ASFV A16/G9/J5 (pE199L) and F9/L1 (pE248R) proteins are both components of the virion inner envelope (26), which has been suggested to fuse with the endosomal membrane of pig cells to release the naked core containing the ASFV genome (23). While the kinetic class of pE248R remains ambivalent, pE199L is clearly a late gene (60). The insect virus *Spodoptera frugiperda* ascovirus 1a encodes 21 proteins that occur in the virion (61), which include its single A16/G9/J5 protein (YP_762409.1) and two F9/L1 proteins (YP_762390.1 and YP_762391.1). In mimivirus, an amoebozoan-infecting GV, its A16/G9/J5 protein (R557; YP_003987072.1) was detected in the virion proteome (62), and its F9/L1 gene (L323; YP_003986826.1) has a late gene expression promoter element (57) annotated in its genome (NC_014649.1). In the closely related sambavirus, the A16/G9/J5 protein (AMK61942.1) was also detected inside the virion (24). In medusavirus, which also infects the amoebozoan *Acanthamoeba*, the F9/L1 gene (BBI30469.1) and two A16/G9/J5 genes (BBI30243.1 and BBI30262.1) are in gene cluster 5 (the latest class) and the other two A16/G9/J5 genes (BBI30251.1 and BBI30253.1) are in cluster 4 (the second latest class) (63). To summarize, for both *Poxviridae* and other GVs that infect animals or amoebozoans, their A16/G9/J5 and F9/L1 genes tend to have late expression and their protein products occur in the virion proteome.

## Discussion

The EFC proteins A16/G9/J5 and F9/L1 have been found to be essential and non-redundant for cell entry, in particular the membrane fusion step, by VacV (11) and ASFV (25, 26), which are the prototypical viruses of *Poxviridae* and *Asfarviridae* that comprise the class *Pokkesviricetes* of *Nucleocytoviricota*. Both A16/G9/J5 and F9/L1 families are cysteine-rich proteins with transmembrane domains near the C-terminal end (11). In this study we show that these two generally co-occurring protein families are commonly distributed in both classes and most major lineages of *Nucleocytoviricota*. The overall conservation in the gene sequences and the predicted proteins suggests that these EFC homologs likely have a conserved function in the virus entry-fusion process across a wide range of GVs similar to that in VacV and ASFV.

One important implication for GV biology is that membrane fusion mediated by A16/G9/J5 and F9/L1 proteins is possibly a common mechanism for cell entry. Fusion with endosome or plasma membrane has been observed for many GVs infecting animal or amoebozoan cells. The absence of A16/G9/J5 and F9/L1 genes in algal GVs, from the families *Phycodnaviridae* and *Mimiviridae*, might be related to the presence of cell walls in most algae, which are an extra barrier that requires more sophisticated strategies for viral entry. For example, when the chlorovirus virion enters the *Chlorella* cell, its spike structure and enzymes first puncture the cell wall, before the virion internal membrane fuses with the cell membrane to form a tunnel for delivering viral DNA into the cytoplasm (19). However, A16/G9/J5 and F9/L1 are also absent in prasinovirus that infects wall-less green algal picoplankton (64), phaeovirus that infects wall-less gametes or spores of brown algae (65), and coccolithovirus infecting coccolithophores using an animal-like strategy (66). Although algal GVs tend to have smaller genome size than the amoeba-infecting GVs (13), we note that the distribution of A16/G9/J5 and F9/L1 genes does not correlate with gene repertoire size, which is even smaller in poxviruses and iridoviruses that mostly have these EFC genes. Instead, a more plausible explanation for the general absence of A16/G9/J5 and F9/L1 in GVs isolated from algae is that the genes were already lost in their ancient ancestors. Based on a recent taxonomic treatment (48), the most parsimonious scenario would be complete losses of both A16/G9/J5 and F9/L1 at the origins of three lineages: 1) a subclade of *Imitervirales*, including algal viruses such as PgV, CpV (both within mesomimiviridae), TetV, and AaV, and ChoanoV infecting a choanoflagellate, 2) *Algavirales*, including chloroviruses, prasinoviruses, and other algal viruses, and 3) pandoravirales (or its major subclade), including coccolithoviruses, phaeoviruses, and amoeba-infecting pandoraviruses and mollivirus where EFC genes are absent. Some smaller subclades within Mimiviridae might also have independently lost the EFC genes, such as those containing the viruses infecting the non-amoebozoan protists, *Bodo* (Discoba) and *Cafeteria* (Stramenopila). Interestingly, all GVs known to infect algae and non-amoebozoan heterotrophic microeukaryotes do not have A16/G9/J5 and F9/L1 homologs, suggesting that the loss of these entry-fusion proteins that work well in animal and amoebozoan hosts might have been the driving force for their ancestors to adapt to other hosts.

The most widely distributed genes among GVs are known to encode proteins including DNA polymerase elongation subunit family B, D5-like primase-helicase, proliferating cell nuclear antigen, DNA-directed RNA polymerase subunits alpha and beta, poxvirus late transcription factor VLTF3, major capsid protein, and packaging ATPase (48). Although A16/G9/J5 and F9/L1 are not as commonly found in individual GVs, their distribution (Figure 1) and phylogenetic (Figures 2 and 3) patterns suggest their presence in LNCA. This adds to the complexity of LNCA, which, in addition to the aforementioned proteins related to DNA replication, RNA transcription, and virion assembly, possessed membrane fusion proteins for virus entry through the endocytic or plasma membrane pathway. It also implies that the host cells of LNCA were more similar to present-day cells of animals (Opisthokonta) and Amoebozoa, which both belong to one of the two major clades of eukaryotes, Amorphea (67). This agrees with the hypothesis that LNCA originated after the origin of eukaryotic cells (68) and the observation that *Nucleocytoviricota* viruses are only known to infect eukaryotes. Given the results presented in this study, A16/G9/J5 and F9/L1 are likely innovations of LNCA or its ancestors. We speculate that these genes for virus entry, along with those for DNA replication, RNA transcription, and virion assembly, could have paved the way for the descendants of LNCA to infect diverse animal (vertebrate and invertebrate) and amoebozoan (Discosea and Tubulinea) hosts and set the stage for later genome gigantism.

To date, mechanisms of viral membrane fusion have only been proposed for the fusion process induced by enveloped viruses carrying only one fusion protein (9). How poxviruses use a 11-protein complex to mediate membrane fusion remains a mystery. Nevertheless, the basic principles of membrane fusion are likely similar, such as the presence of a hydrophobic fusion loop or fusion peptide for insertion into host membrane and the conformational change that breaks the energy barrier and ultimately merges two membranes. Accordingly, if A16/G9/J5 and F9/L1 were the components in ancestors of GVs that worked together to drive membrane fusion, either one of them should contain the fusion loop or fusion peptide. Foo et al. (69) proposed a myristoyl switch model for VacV L1 in which the buried N-terminal myristate moiety in the L1 helix α1 swings up upon activation and the exposed N-terminal lipid group acts as an anchor to connect the host membrane. Interestingly, although both poxviral A16 and G9 harbor an N-myristoylation motif “MG”, this lipid modification seems more conserved in the F9/L1 family of GVs (Figures 2 and 3). Besides, a recent study showed that abrogation of VacV L1 N-myristoylation is more detrimental to the virus than inhibiting the same modification on A16 or G9 (54). Further studies of L1 are warranted to unveil the function of L1 N-myristoylation and its role in GV entry. Also of note, the presence of A16/G9/J5 homologs in many GVs raises the question of whether proteins similar to the VacV fusion suppressers, namely A26 (70) and A56/K2 complex (71), that interact with VacV A16/G9 subcomplex to regulate fusion, also exist in other GVs.

In this study, we aimed to extend homology search from sequence to structural conservation. We therefore employed AlphaFold2 to predict 3-dimensional structures of selected EFC protein homologs and compared structural similarity by superposition. Strikingly, after selecting models with high pLDDT scores, we found remarkable structural homology in both A16/G9/J5 and F9/L1 families between VacV and distantly related GVs, such as klosneuvirus and tupanvirus deep ocean (Figures 5 and 6). As mentioned above, both A16/G9/J5 and F9/L1 families comprise cysteine-rich proteins and the putative disulfide bonding patterns appear conserved across GVs (Figures 2 and 3). Therefore, we deduced that the intramolecular disulfide bonding may be associated with an evolutionarily conserved entry-fusion mechanism for GV entry.

In addition to implications for protein conservation, host range, and ancient evolution, this study highlights A16/G9/J5 and F9/L1 proteins in GVs as targets for further biochemical and molecular biological characterization. Unlike other GVs, poxviruses are highly conserved in having three A16/G9/J5 and two F9/L1 genes (Figure 1). In addition to individual protein structures (Figures 5 and 6), their structural and functional roles in the entire poxvirus EFC remain to be elucidated. The general co-occurrence of A16/G9/J5 and F9/L1 genes in other GVs (Figure 1) also suggests their protein products might interact and coevolve with each other. Some intriguing examples of close proximity of A16/G9/J5 and F9/L1 genes and their conserved genomic localizations (Figure 4) merit further investigation of their roles in expression and functional regulation. With the importance of GVs for public health, agriculture, wildlife populations, and environmental microbes, a comprehensive view of their entry-fusion proteins is greatly needed and pivotal for understanding their biology and evolution.

## Supplementary Material

Supplemental material is available online.

Table S1: List of A16/G9/J5 and F9/L1 protein homologs identified in this study.

Table S2: List of EFC homologs in carp edema virus strain FTI2020, saltwater crocodilepox virus subtype 1, teiidaepox virus 1, and cheloniid poxvirus 1.

Data S1: Maximum likelihood tree of the A16/G9/J5 gene family.

Data S2: Maximum likelihood tree of the F9/L1 gene family.

Data S3: Structural models of A16/G9/J5 and F9/L1 proteins from selected viruses.

## Acknowledgments

This work was funded by Academia Sinica Career Development Award, grant number AS-CDA-110-L01 (C.K.) and National Science and Technology Council, Taiwan, grant numbers 108-2311-B-001-040-MY3 and 111-2611-M-001-008-MY3 (C.K.) and 110-2320-B-001-015-MY3 (W.C.). The authors thank Tsu-Wang Sun, Tzu-Tong Kao, the laboratory of Chih-Horng Kuo, and Hsin-Nan Lin (Bioinformatics Core, Institute of Molecular Biology) for assistance with bioinformatic analyses.

## Notes

### Competing Interest Statement

The authors have declared no competing interest.

## References

1. Chung C-S, Hsiao J-C, Chang Y-S, Chang W. 1998. A27L Protein Mediates Vaccinia Virus Interaction with Cell Surface Heparan Sulfate. J Virol 72:1577–1585.

2. Lin C-L, Chung C-S, Heine HG, Chang W. 2000. Vaccinia Virus Envelope H3L Protein Binds to Cell Surface Heparan Sulfate and Is Important for Intracellular Mature Virion Morphogenesis and Virus Infection In Vitro and In Vivo. J Virol 74:3353–3365.

3. Chiu W-L, Lin C-L, Yang M-H, Tzou D-LM, Chang W. 2007. Vaccinia Virus 4c (A26L) Protein on Intracellular Mature Virus Binds to the Extracellular Cellular Matrix Laminin. J Virol 81:2149–2157.

4. Izmailyan R, Hsao J-C, Chung C-S, Chen C-H, Hsu PW-C, Liao C-L, Chang W. 2012. Integrin β1 Mediates Vaccinia Virus Entry through Activation of PI3K/Akt Signaling. J Virol 86:6677–6687.

5. Schroeder N, Chung C-S, Chen C-H, Liao C-L, Chang W. 2012. The Lipid Raft-Associated Protein CD98 Is Required for Vaccinia Virus Endocytosis. J Virol 86:4868–4882.

6. Huang C-Y, Lu T-Y, Bair C-H, Chang Y-S, Jwo J-K, Chang W. 2008. A Novel Cellular Protein, VPEF, Facilitates Vaccinia Virus Penetration into HeLa Cells through Fluid Phase Endocytosis. J Virol 82:7988–7999.

7. Hsiao J-C, Chu L-W, Lo Y-T, Lee S-P, Chen T-J, Huang C-Y, Ping Y-H, Chang W. 2015. Intracellular Transport of Vaccinia Virus in HeLa Cells Requires WASH-VPEF/FAM21-Retromer Complexes and Recycling Molecules Rab11 and Rab22. J Virol 89:8365–8382.

8. Townsley AC, Weisberg AS, Wagenaar TR, Moss B. 2006. Vaccinia Virus Entry into Cells via a Low-pH-Dependent Endosomal Pathway. J Virol 80:8899–8908.

9. Harrison SC. 2015. Viral membrane fusion. Virology 479–480:498–507.

10. Moss B. 2012. Poxvirus cell entry: How many proteins does it take? Viruses 4:688–707.

11. Moss B. 2016. Membrane fusion during poxvirus entry. Semin Cell Dev Biol 60:89–96.

12. Koonin E V., Dolja V V., Krupovic M, Varsani A, Wolf YI, Yutin N, Zerbini FM, Kuhn JH. 2020. Global Organization and Proposed Megataxonomy of the Virus World. Microbiol Mol Biol Rev 84:e00061–19.

13. Sun T-W, Yang C-L, Kao T-T, Wang T-H, Lai M-W, Ku C. 2020. Host Range and Coding Potential of Eukaryotic Giant Viruses. Viruses 12:1337.

14. Van Etten JL, Meints RH. 1999. Giant viruses infecting algae. Annu Rev Microbiol 53:447–94.

15. Claverie J-M, Abergel C. 2016. Giant viruses: The difficult breaking of multiple epistemological barriers. Stud Hist Philos Biol Biomed Sci 59:89–99.

16. Ku C. 2021. Giant Virus-Eukaryote Interactions as Ecological and Evolutionary Driving Forces. mSystems 6:e00737–21.

17. Schulz F, Abergel C, Woyke T. 2022. Giant virus biology and diversity in the era of genome-resolved metagenomics. Nat Rev Microbiol. Springer US.

18. Sun T-W, Ku C. 2021. Unraveling Gene Content Variation Across Eukaryotic Giant Viruses Based on Network Analyses and Host Associations. Virus Evol 7:veab081.

19. Van Etten JL, Agarkova I V, Dunigan DD. 2020. Chloroviruses. Viruses 12:20.

20. Abergel C, Legendre M, Claverie JM. 2015. The rapidly expanding universe of giant viruses: Mimivirus, Pandoravirus, Pithovirus and Mollivirus. FEMS Microbiol Rev 39:779–796.

21. de Souza GAP, Queiroz VF, Coelho LFL, Abrahão JS. 2021. Alohomora! What the entry mechanisms tell us about the evolution and diversification of giant viruses and their hosts. Curr Opin Virol 47:79–85.

22. Wang S, Huang X, Huang Y, Hao X, Xu H, Cai M, Wang H, Qin Q. 2014. Entry of a Novel Marine DNA Virus, Singapore Grouper Iridovirus, into Host Cells Occurs via Clathrin-Mediated Endocytosis and Macropinocytosis in a pH-Dependent Manner. J Virol 88:13047–13063.

23. Andrés G. 2017. African Swine Fever Virus Gets Undressed: New Insights on the Entry Pathway. J Virol 91:1–5.

24. Schrad JR, Abrahão JS, Cortines JR, Parent KN. 2020. Structural and Proteomic Characterization of the Initiation of Giant Virus Infection. Cell 181:1046–1061.e6.

25. Hernáez B, Guerra M, Salas ML, Andrés G. 2016. African Swine Fever Virus Undergoes Outer Envelope Disruption, Capsid Disassembly and Inner Envelope Fusion before Core Release from Multivesicular Endosomes. PLoS Pathog 12:1–32.

26. Matamoros T, Alejo A, Rodríguez JM, Hernáez B, Guerra M, Fraile-Ramos A, Andrés G. 2020. African swine fever virus protein pE199L mediates virus entry by enabling membrane fusion and core penetration. MBio 11:1–21.

27. Yutin N, Koonin E V. 2012. Hidden evolutionary complexity of Nucleo-Cytoplasmic Large DNA viruses of eukaryotes. Virol J 9:1–18.

28. Moniruzzaman M, Martinez-Gutierrez CA, Weinheimer AR, Aylward FO. 2020. Dynamic genome evolution and blueprint of complex virocell metabolism in globally-distributed giant viruses. Nat Commun 11:1710.

29. Schulz F, Roux S, Paez-Espino D, Jungbluth S, Walsh DA, Denef VJ, McMahon KD, Konstantinidis KT, Eloe-Fadrosh EA, Kyrpides NC, Woyke T. 2020. Giant virus diversity and host interactions through global metagenomics. Nature 578:432–436.

30. Mekata T, Kawato Y, Ito T. 2021. Complete Genome Sequence of Carp Edema Virus Isolated from Koi Carp. Microbiol Resour Announc 10:8–9.

31. Sarker S, Isberg SR, Milic NL, Lock P, Helbig KJ. 2018. Molecular characterization of the first saltwater crocodilepox virus genome sequences from the world’s largest living member of the Crocodylia. Sci Rep 8:1–11.

32. Seitz K, Kübber-Heiss A, Auer A, Dinhopl N, Posautz A, Mötz M, Kiesler A, Hochleithner C, Hochleithner M, Springler G, Lehmbecker A, Weissenböck H, Rümenapf T, Riedel C. 2021. Discovery of a phylogenetically distinct poxvirus in diseased Crocodilurus amazonicus (family Teiidae). Arch Virol 166:1183–1191.

33. Sarker S, Hannon C, Athukorala A, Bielefeldt-Ohmann H. 2021. Emergence of a novel pathogenic poxvirus infection in the endangered green sea turtle (Chelonia mydas) highlights a key threatening process. Viruses 13.

34. Altschul SF, Madden TL, Schäffer AA, Zhang J, Zhang Z, Miller W, Lipman DJ. 1997. Gapped BLAST and PSI-BLAST: A new generation of protein database search programs. Nucleic Acids Res 25:3389–3402.

35. Yutin N, Wolf YI, Raoult D, Koonin E V. 2009. Eukaryotic large nucleo-cytoplasmic DNA viruses: clusters of orthologous genes and reconstruction of viral genome evolution. Virol J 6:223.

36. Benson DA, Karsch-Mizrachi I, Clark K, Lipman DJ, Ostell J, Sayers EW. 2012. GenBank. Nucleic Acids Res 40:D48–53.

37. Katoh K, Standley DM. 2013. MAFFT multiple sequence alignment software version 7: improvements in performance and usability. Mol Biol Evol 30:772–80.

38. Minh BQ, Schmidt HA, Chernomor O, Schrempf D, Woodhams MD, von Haeseler A, Lanfear R. 2020. IQ-TREE 2: New Models and Efficient Methods for Phylogenetic Inference in the Genomic Era. Mol Biol Evol 37:1530–1534.

39. Kalyaanamoorthy S, Minh BQ, Wong TKF, Von Haeseler A, Jermiin LS. 2017. ModelFinder: Fast model selection for accurate phylogenetic estimates. Nat Methods 14:587–589.

40. Yu G, Smith DK, Zhu H, Guan Y, Lam TTY. 2017. ggtree: an R Package for Visualization and Annotation of Phylogenetic Trees With Their Covariates and Other Associated Data. Methods Ecol Evol 8:28–36.

41. Diesterbeck US, Gittis AG, Garboczi DN, Moss B. 2018. The 2.1 Å structure of protein F9 and its comparison to L1, two components of the conserved poxvirus entry-fusion complex. Sci Rep 8:1–12.

42. Su HP, Garman SC, Allison TJ, Fogg C, Moss B, Garboczi DN. 2005. The 1.51-Å structure of the poxvirus L1 protein, a target of potent neutralizing antibodies. Proc Natl Acad Sci U S A 102:4240–4245.

43. Jumper J, Evans R, Pritzel A, Green T, Figurnov M, Ronneberger O, Tunyasuvunakool K, Bates R, Žídek A, Potapenko A, Bridgland A, Meyer C, Kohl SAA, Ballard AJ, Cowie A, Romera-Paredes B, Nikolov S, Jain R, Adler J, Back T, Petersen S, Reiman D, Clancy E, Zielinski M, Steinegger M, Pacholska M, Berghammer T, Bodenstein S, Silver D, Vinyals O, Senior AW, Kavukcuoglu K, Kohli P, Hassabis D. 2021. Highly accurate protein structure prediction with AlphaFold. Nature 596:583–589.

44. Mirdita M, Schütze K, Moriwaki Y, Heo L, Ovchinnikov S, Steinegger M. 2022. ColabFold: making protein folding accessible to all. Nat Methods 19:679–682.

45. Steinegger M, Söding J. 2017. MMseqs2 enables sensitive protein sequence searching for the analysis of massive data sets. Nat Biotechnol.

46. Schrödinger. 2015. The PyMOL Molecular Graphics Suystem. 1.8.

47. Krissinel E, Henrick K. 2004. Secondary-structure matching (SSM), a new tool for fast protein structure alignment in three dimensions. Acta Crystallogr Sect D Biol Crystallogr 60:2256–2268.

48. Aylward FO, Moniruzzaman M, Ha AD, Koonin E V. 2021. A phylogenomic framework for charting the diversity and evolution of giant viruses. PLOS Biol 19:e3001430.

49. Subramaniam K, Behringer DC, Bojko J, Yutin N, Clark AS, Bateman KS, van Aerle R, Bass D, Kerr RC, Koonin E V., Stentiford GD, Waltzek TB. 2020. A new family of DNA viruses causing disease in crustaceans from diverse aquatic biomes. MBio 11.

50. Yoshikawa G, Blanc-Mathieu R, Song C, Kayama Y, Mochizuki T, Murata K, Ogata H, Takemura M. 2019. Medusavirus, a Novel Large DNA Virus Discovered from Hot Spring Water. J Virol 93:e02130–18.

51. Tully BJ, Wheat CG, Glazer BT, Huber JA. 2018. A dynamic microbial community with high functional redundancy inhabits the cold, oxic subseafloor aquifer. ISME J 12:1–16.

52. Takahashi H, Fukaya S, Song C, Murata K, Takemura M. 2021. Morphological and Taxonomic Properties of the Newly Isolated Cotonvirus japonicus, a New Lineage of the Subfamily Megavirinae. J Virol 95.

53. Ojeda S, Domi A, Moss B. 2006. Vaccinia Virus G9 Protein Is an Essential Component of the Poxvirus Entry-Fusion Complex. J Virol 80:9822–9830.

54. Priyamvada L, Kallemeijn WW, Faronato M, Wilkins K, S. Cg, Cotter CA, Ojeda S, Solar R, Moss BW, Panayampalli T, Satheshkumar S. 2022. Inhibition of vaccinia virus L1 N-myristoylation by the host N-myristoyltransferase inhibitor IMP-1088 generates non-infectious virions defective in cell entry. PLOS Pathog 18:e1010662.

55. Martin KH, Grosenbach DW, Franke CA, Hruby DE. 1997. Identification and analysis of three myristylated vaccinia virus late proteins. J Virol 71:5218–5226.

56. Ku C, Sheyn U, Sebé-Pedrós A, Ben-Dor S, Schatz D, Tanay A, Rosenwasser S, Vardi A. 2020. A single-cell view on alga-virus interactions reveals sequential transcriptional programs and infection states. Sci Adv 6:eaba4137.

57. Legendre M, Audic S, Poirot O, Hingamp P, Seltzer V, Byrne D, Lartigue A, Lescot M, Bernadac A, Poulain J, Abergel C, Claverie JM. 2010. mRNA deep sequencing reveals 75 new genes and a complex transcriptional landscape in Mimivirus. Genome Res 20:664–674.

58. Oliveira GP, Andrade AC dos SP, Rodrigues RAL, Arantes TS, Boratto PVM, Silva LKDS, Dornas FP, Trindade G de S, Drumond BP, La Scola B, Kroon EG, Abrahï¿½o JS. 2017. Promoter motifs in NCLDVs: An evolutionary perspective. Viruses 9:1–20.

59. Assarsson E, Greenbaum JA, Sundström M, Schaffer L, Hammond JA, Pasquetto V, Oseroff C, Hendrickson RC, Lefkowitz EJ, Tscharke DC, Sidney J, Grey HM, Head SR, Peters B, Sette A. 2008. Kinetic analysis of a complete poxvirus transcriptome reveals an immediate-early class of genes. Proc Natl Acad Sci U S A 105:2140–5.

60. Cackett G, Matelska D, Sýkora M, Portugal R, Malecki M, Bähler J, Dixon L, Werner F. 2020. The African swine fever virus transcriptome. J Virol 94:1–22.

61. Tan Y, Bideshi DK, Johnson JJ, Bigot Y, Federici BA. 2009. Proteomic analysis of the Spodoptera frugiperda ascovirus 1a virion reveals 21 proteins. J Gen Virol 90:359–365.

62. Renesto P, Abergel C, Decloquement P, Moinier D, Azza S, Ogata H, Fourquet P, Gorvel J-P, Claverie J-M. 2006. Mimivirus Giant Particles Incorporate a Large Fraction of Anonymous and Unique Gene Products. J Virol 80:11678–11685.

63. Zhang R, Endo H, Takemura M, Ogata H. 2021. RNA Sequencing of Medusavirus Suggests Remodeling of the Host Nuclear Environment at an Early Infection Stage. Microbiol Spectr 9.

64. Derelle E, Ferraz C, Escande ML, Eychenié S, Cooke R, Piganeau G, Desdevises Y, Bellec L, Moreau H, Grimsley N. 2008. Life-cycle and genome of OtV5, a Large DNA virus of the pelagic marine unicellular green alga Ostreococcus tauri. PLoS One 3.

65. Van Etten JL, Lane LC, Dunigan DD. 2010. DNA Viruses: The Really Big Ones (Giruses). Annu Rev Microbiol 64:83–99.

66. Mackinder LCM, Worthy CA, Biggi G, Hall M, Ryan KP, Varsani A, Harper GM, Wilson WH, Brownlee C, Schroeder DC. 2009. A unicellular algal virus, Emiliania huxleyi virus 86, exploits an animal-like infection strategy. J Gen Virol 90:2306–2316.

67. Adl SM, Bass D, Lane CE, Lukeš J, Schoch CL, Smirnov A, Agatha S, Berney C, Brown MW, Burki F, Cárdenas P, Čepička I, Chistyakova L, del Campo J, Dunthorn M, Edvardsen B, Eglit Y, Guillou L, Hampl V, Heiss AA, Hoppenrath M, James TY, Karpov S, Kim E, Kolisko M, Kudryavtsev A, Lahr DJG, Lara E, Le Gall L, Lynn DH, Mann DG, Massana i Molera R, Mitchell EAD, Morrow C, Park JS, Pawlowski JW, Powell MJ, Richter DJ, Rueckert S, Shadwick L, Shimano S, Spiegel FW, Torruella i Cortes G, Youssef N, Zlatogursky V, Zhang Q, Zhang Q. 2019. Revisions to the classification, nomenclature, and diversity of eukaryotes. J Eukaryot Microbiol 66:4–119.

68. Ku C, Sun T. 2020. Did giant and large dsDNA viruses originate before their eukaryotic hosts? Proc Natl Acad Sci 117:2747–2748.

69. Foo CH, Whitbeck JC, Ponce-de-León M, Saw WT, Cohen GH, Eisenberg RJ. 2012. The Myristate Moiety and Amino Terminus of Vaccinia Virus L1 Constitute a Bipartite Functional Region Needed for Entry. J Virol 86:5437–5451.

70. Chang S-J, Shih A-C, Tang Y-L, Chang W. 2012. Vaccinia Mature Virus Fusion Regulator A26 Protein Binds to A16 and G9 Proteins of the Viral Entry Fusion Complex and Dissociates from Mature Virions at Low pH. J Virol 86:3809–3818.

71. Wagenaar TR, Ojeda S, Moss B. 2008. Vaccinia Virus A56/K2 Fusion Regulatory Protein Interacts with the A16 and G9 Subunits of the Entry Fusion Complex. J Virol 82:5153–5160.

